# Cross-seeding Controls Aβ Fibril Populations and Resulting Function

**DOI:** 10.1101/2021.10.14.464427

**Authors:** Michael J. Lucas, Henry S. Pan, Eric J. Verbeke, Gina M. Partipilo, Ethan C. Helfman, Leah Kann, Benjamin K. Keitz, David W. Taylor, Lauren J. Webb

## Abstract

Amyloid peptides nucleate from monomers to aggregate into fibrils through primary nucleation; pre-existing fibrils can then act as seeds for additional monomers to fibrillize through secondary nucleation. Both nucleation processes can occur simultaneously, yielding a distribution of fibril polymorphs that can generate a spectrum of neurodegenerative effects. Understanding the mechanisms driving polymorph structural distribution during both nucleation processes is important for uncovering fibril structure-function relationships, as well creating polymorph distributions *in vitro* that better match distributions found *in vivo*. Here, we explore how cross-seeding WT Aβ_1-40_ with Aβ_1-40_ mutants E22G (Arctic) and E22Δ (Osaka), as well as with WT Aβ_1-42_ affects the distribution of fibril structural polymorphs, and how changes in structural distribution impact toxicity. Transmission electron microscopy analysis reveals that fibril seeds derived from mutants of Aβ_1-40_ impart their structure to WT Aβ_1-40_ monomer during secondary nucleation, but WT Aβ_1-40_ fibril seeds do not affect the structure of fibrils assembled from mutant Aβ_1-40_ monomers, despite kinetics data indicating accelerated aggregation when cross-seeding of any combination of mutants. Additionally, WT Aβ_1-40_ fibrils seeded with mutant fibrils to produce similar structural distributions to the mutant seeds also produced similar cytotoxicity on neuroblastoma cell lines. This indicates that mutant fibril seeds not only impart their structure to growing WT Aβ_1-40_ aggregates, but they also impart cytotoxic properties. Our findings provide clear evidence that there is a relationship between fibril structure and phenotype on a polymorph population level, and that these properties can be passed on through secondary nucleation of succeeding generations of fibrils.

## Introduction

A number of neurodegenerative diseases are characterized by misfolded proteins that nucleate from monomeric peptides into soluble oligomers, ultimately resulting in insoluble fibrils.^1,2^ While oligomers show significant neurotoxicity,^3–6^ the study of amyloid fibrils can provide important insight on the mechanism of amyloid formation in diseases for several reasons. First, since amyloid formation is a nucleation driven process, fibrils can serve as a source of secondary nucleation, further catalyzing the aggregation of monomeric peptide from a structured seed.^7–10^ Second, because fibrils are more stable than their oligomeric counter-parts,^11^ they can be used as easily observed biomarkers that are indicative of the entire aggregation process. Finally, amyloidogenic proteins can form fibrils with distinct structural differences, which are hypothesized to be the result of initial nucleation events.^12^ The ability for fibrils to adopt different structures is known as structural polymorphism.^13^ For example, varying fibrillization conditions can yield a distribution of different polymorphs that vary in width, helicity, and crossover distance—the distance required for the fibril to complete a 180° rotation. Given that fibrils form in a variety of cellular compartments and that changes in the distribution of structural polymorphs can lead to different disease states, the connection between physiological state, structural polymorphism, biochemical properties, and phenotype poses one of the main challenges in understanding the role of fibrils in neurodegeneration.^14–18^

Amyloid fibrils are characterized by a cross-β motif, in which β-sheets extend perpendicular to the fibril axis and stack together to form protofilaments.^19–21^ Differences in the registry and stacking of the β-sheets and protofilament symmetry give rise to structural variants of fibrils formed by the same peptide.^13^ For example, fibrils formed from amyloid-β (Aβ) can contain two or three protofilaments with varying symmetry.^22^ Additionally, differences in packing and symmetry can lead to the formation of multiple potential steric-zippers, each of which are able to serve as the spine of the fibril.^13^ Because fibrils are polymorphic, the number of conformations available to a single peptide sequence can be extremely large.^13^ As recent studies have shown that structural variants can display differing phenotypes and disease subtypes,^23–27^ the large number of possible fibril conformations complicates the development of effective therapeutics and hinders our understanding of the role of fibrils in disease progression.

The pathophysiology of Alzheimer’s Disease (AD) is attributed to two amyloidogenic peptides: Aβ and tau.^28^ A number of *in vitro* structures have been identified for both Aβ and tau fibrils using NMR spectroscopy, X-ray crystallography, and transmission electron microscopy (TEM).^29–31^ These studies have demonstrated that incubation conditions – including temperature, shaking, salt concentration and pH – can influence fibril structure. In addition to the fibril polymorphs identified *in vitro*, recent studies have elucidated the structure of several fibril polymorphs from patient-derived brains *ex vivo*.^32,33^ Unfortunately, there are several structural differences between Aβ and tau fibrils prepared *in vitro* and their *ex vivo* counterparts, where peptide conformation at the fibril core and even the handedness of the fibrils themselves differ. These differences are most likely due to post-translation modifications, amino acid sequence variation between isoforms, and cellular microenvironments to which the peptides are exposed.^34^ A small portion of AD cases are associated with familial mutations (FAD) in the amyloid precursor protein, which have been shown to result in significant differences in structural polymorphism of the resulting Aβ fibrils. For example, a single-residue change within positions 21-23 of Aβ removes a potential salt bridge and results in a wide variety of fibril structures that differ in width and crossover distance.^35–37^ There is also increasing evidence that these mutations lead to a diverse array of disease phenotypes. In mouse models, fibrils formed from the Osaka mutation (E22Δ) had accelerated aggregation and gained increased toxicity when compared to wild-type (WT) Aβ.^38^ Additionally, recent studies have shown that mutations of residues 21-23 of Aβ can affect the kinetics of nucleation since the resulting fibrils display differences in surface hydrophobicity, potentially altering protein-protein interactions.^39^ Overall, the numerous fibril structures formed by familial mutants and wild-type (WT) Aβ further complicates our understanding of the pathological role of fibrils and their structural polymorphism.

Recent studies have revealed that cross-seeding occurs between a variety of amyloidogenic proteins during disease progression.^40^ The cross-seeding of wild-type Aβ_1-40_ with various FAD mutants and isoforms can yield multiple fibril polymorphs that are different from the parent peptide and that influence the formation of additional structures, as well as cause changes in phenotype. Seeding any aggregating peptide monomer with pre-formed fibrils is a common method for producing a new generation of fibrils as microscopic structure propagates from parent to progeny to create homogenous populations.^41–43^ Previous studies have shown that when seeding amyloid fibrils, the phenotype of the fibril population can change after multiple generations, leading to more interest in establishing structure-function relationships.^44^ Cross-seeding between different amyloid isoforms and mutants *in vitro* has also gained attention in recent years to yield insights into the complex mechanism of aggregation *in vivo*, including studies in which Aβ is subjected to various microenvironments and post-translational modifications. However, a majority of these studies focused on the kinetics of fibrilization rather than the structural and phenotypic impact, and there is a large library of potentially relevant fibril structures.^45,46^ Cross-seeding between different forms of Aβ has the potential to contribute to increased fibril polymorph diversity, which can likely lead to different pathological outcomes. For example, previous studies have demonstrated that heterozygous carriers of the E22Δ mutation (Osaka) in murine models do not display signs of dementia, while those heterozygous with the E22G mutation (Arctic) do show significant dementia.^47,48^ By understanding fibril propagation in these diseases and how propagation affects the distribution of fibril structural polymorphs, our goal is to better understand the phenotypes that are observed in these diseases.

In this study, we use cross-seeding methods, specifically focusing on Aβ, to attempt to better understand the propagation of fibrils and their resulting structural polymorphism. Using electron microscopy, we conducted experiments to measure the distribution of fibril structural polymorphs produced from individual monomer sequences, and cross-seeded sequences. We examined the structural and phenotypic effects of multiple generations of seeding WT Aβ_1-40_ as well as cross-seeding with FAD mutants or WT Aβ_1-42_. We evaluated the impact of seeding WT Aβ_1-40_ with pre-formed fibrils of the Aβ Arctic and Osaka mutants. From these different seeding conditions, we determined the structural polymorphism distribution of the resulting fibrils with TEM and the resulting impact on human neuroblastoma cell viability. Our results demonstrate that after multiple generations of Aβ_1-40_ seeding, a homogenous population of fibrils with a crossover distance of 30 nm was formed, which resulted in decreased cell viability when compared to fibrils with a longer crossover distance. Additionally, we show that structure can be reproducibly passed from mutant fibrils to that of WT fibrils. Cross-seeding of WT fibrils resulted in a toxicity profile similar to that of the parent mutant fibrils, demonstrated by cellular assays. Our findings have important implications for neurodegenerative diseases caused by the aggregation of monomers into fibrils, and suggest that distinct fibril populations, not just specific structures, may alter disease pathology.

## Results

We chose to focus our studies on the Arctic (E22G) and Osaka (E22Δ) mutations (Figure 1a) because of the importance mutations at the Glu at position 22 of Aβ. In the previously solved structures of Aβ_1-40_ fibrils, E22 is exposed to the solvent, allowing it to interact with the surrounding environment, including other Aβ monomers.^30^ Additionally, since Glu is negatively charged, E22G and E22Δ result in a one unit change in net charge at the exposed fibril surface. Since electrostatics, sterics, and the repulsion between the negatively charged Aβ peptides are likely important in the aggregation mechanism, a change in charge could significantly change aggregation behavior.^49^ Furthermore, in some structures of Aβ_1-40_, residue D23 forms a salt bridge with K28.^41^ However, a NMR characterized structure of the E22Δ mutation showed that in the absence of E22, D23 becomes solvent exposed, both removing the salt bridge with K28 and changing peptide conformation at the fibril core.^50^ Because secondary nucleation occurs on fibril surfaces, these changes to fibril structure can impact the seeding capabilities these mutant fibrils have in the presence of additional Aβ_1-40_ monomer. Finally, the E22 mutations lead to distinct disease phenotypes; the E22G mutation enhances protofibril formation and proteolysis-resistant aggregates, while the E22Δ mutation increases intracellular aggregation.^51^ Given the structural importance of E22, the distinct disease phenotypes, and lack of characterized structures for these mutations, we focused our efforts on understanding the structure of Arctic and Osaka mutations and their seeding capabilities with WT Aβ_1-40_. In addition to the cross-seeding interactions between the mutant forms of Aβ with WT Aβ_1-40_, we also examined how the inclusion of the more toxic WT Aβ_1-42_ isoform would affect cross-seeding interactions, structural polymorphic distributions, and resulting phenotypes by cellular assays (Figure 1b).

**Figure 1.**
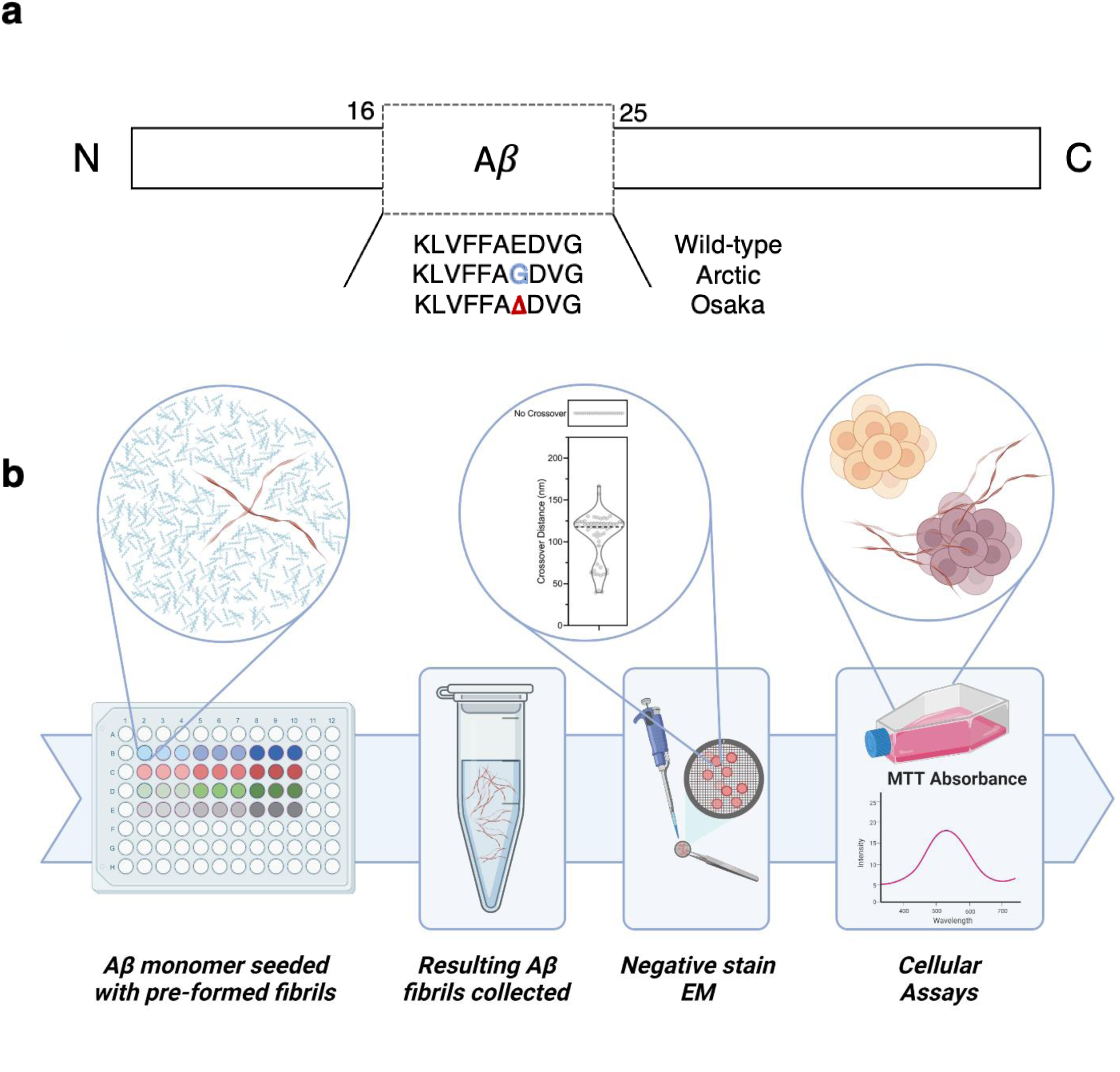
a) Sequence of residues 16-25 of wild-type, Arctic (E22G), and Osaka (E22Δ) Aβ. b) Workflow of the preparation and characterization of fibril polymorphs. 30 μM Aβ_1-40_ and its mutants were incubated with 5 μM of pre-formed fibrils for 24 h at 37 °C. Prepared fibrils were independently imaged by negative stain TEM to characterize structural polymorphs and analyzed for their effects on cell viability.

### Structural Analysis of Mutant Fibrils and Their Seeds

To analyze fibril structures, we used TEM to determine helical crossover distances. Our initial studies examined repeated seeding, the interactions between different Aβ aggregates, and how these different aggregation conditions influenced fibril helical crossover distance distributions. Previous studies have shown that helical crossover distances are a relatively straightforward method for differentiating different amyloid polymorphs, even though distinct structures could potentially have the same crossover distance.^32^ Compared to other structural characterization methods such as NMR spectroscopy and X-ray crystallography, TEM offers the advantage of examining a wide field of fibrils that do not need to be purified to homogeneity. Therefore, using negative stain TEM, we were able to rapidly characterize and quantify complex populations of fibril polymorphs that formed under various conditions.

After incubating 30 μM of WT Aβ_1-40_ at 37 °C for 24 hours, amyloid fibrils were recovered by centrifugation, drop-cast onto TEM grids, and stained with uranyl acetate. At least 50 images were collected by TEM and analyzed with ImageJ^52^ for each replicate preparation of fibrils to determine helical crossover distance. We then used RELION^53^ to compute reference-free 2D class averages as a way to verify consistency between different morphologies across replicates by sorting images of fibrils into self-similar classes. Figure 2a displays the various crossover distances identified from the class averages. Helical crossover distances varied in length from 30 nm to 120 nm between each crossover point. Additionally, there was a population of fibrils with no crossover distance. As reported previously^22^ and as seen in Figure 2b (Generation 0), WT Aβ_1-40_ primarily formed fibrils with crossover distances of 120 nm (73 ± 14%), 60 nm (7 ± 7%), and no crossovers (20 ± 9%).^54^ In order to evaluate if any fibril polymorphs were able to pass on their structure, we seeded 30 μM of fresh WT Aβ_1-40_ monomer with 5 μM of pre-formed fibril “seeds.” We repeated this process twice to produce 3 generations of fibrils. As seen in Figure 2b, successive generations of seeding fresh WT Aβ_1-40_ monomer with previously formed fibrils led to a homogenous population of fibrils with a crossover distance of 30 nm. This confirmed that fibril structure can be replicated and controlled with pre-formed aggregates, and also demonstrates that fibrils seeded through secondary nucleation preferred to nucleate into a structure of primarily one crossover distance, consistent with previous literature on generational seeding of fibrils to obtain homogeneous samples.^22^

**Figure 2.**
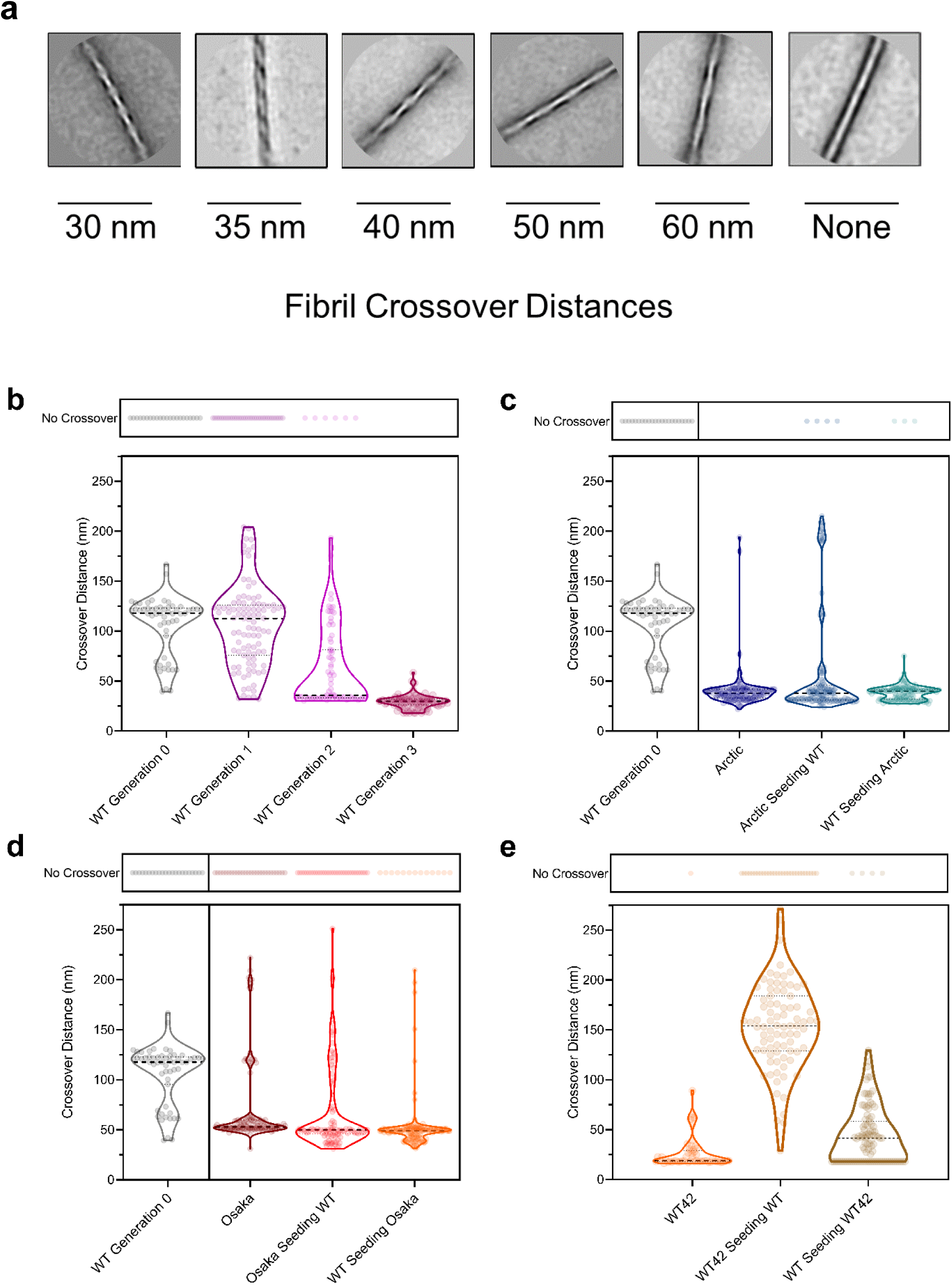
(a) Class average images in 200 × 200 nm boxes of various Aβ fibril crossover distances observed by negative-stain TEM. Structural distributions of fibril polymorphs as measured by crossover distance, for (b) repeated seeding of 30 μM WT Aβ_1-40_ monomer with 5 μM fibril from the previous generation; (c) 30 μM pure Arctic, 5 μM Arctic seeding 30 μM WT, and 5 μM WT seeding 30 μM Arctic; (d) 30 μM pure Osaka, 5 μM Osaka seeding 30 μM WT, and 5 μM WT seeding 30 μM Osaka; (e) 30 μM WT42, 5 μM WT42 seeding 30 μM WT, and 5 μM WT seeding 30 μM WT42. Fibril crossover distance was determined from a total of 50 micrographs from three biological replicates. The dashed line represents the median of the distribution.

We then examined how cross-seeding between WT Aβ_1-40_ and E22 mutants affected fibril polymorphism. We first determined the distribution of fibril crossover distances of Arctic and Osaka fibrils by analyzing the fibrils formed after a 24 hr incubation at 37 °C. Similar to the WT Aβ_1-40_ generational imaging, three replicates each containing 50 micrographs were analyzed for helical crossover distance. As seen in Figure 2c, the Arctic mutation primarily formed fibrils with crossover distances of 40 nm (63 ± 12%) and 30 nm (28 ± 13%), consistent with previous reports.^37^ Similarly, the Osaka mutation produced fibrils of a shorter crossover distance (Figure 2d), with the predominant fibril having a crossover distance of 50 nm (50 ± 20%). However, fibrils with the Osaka mutation yielded a distribution containing a larger portion of fibrils with longer crossover distances than Arctic, exhibiting crossover distances of 200 nm (5 ± 9%), 120 nm (8 ± 7%), 60 nm (21 ± 10%), and no crossover distance (16 ± 13%).

Next, we analyzed the ability of the mutant fibrils to consistently pass their structure into WT Aβ_1-40_ monomer. After incubating 5 μM of pre-formed mutant fibril seeds with 30 μM of Aβ_1-40_ monomer at 37 °C for 24 hours, the resulting fibrils were drop-cast on TEM grids, as described previously, and 50 micrographs were examined across three replicates. When WT Aβ_1-40_ monomer was seeded by Arctic fibrils, a significantly different population of fibrils was formed when compared to a WT Aβ_1-40_ control (Figure 2c). WT Aβ_1-40_ fibrils seeded by the Arctic mutant primarily formed fibrils with crossover distances of 30 nm (45 ± 5%) and 40 nm (27 ± 11%). This population distribution largely reflected that of Arctic fibrils, indicating that the Arctic mutation faithfully replicates its structure in WT Aβ_1-40_ fibrils. There was also a small population of fibrils with crossover distances of 50 nm (6 ± 2%), 60 nm (6 ± 3%), and 200 nm (12 ± 11%). Similarly, WT Aβ_1-40_ monomer seeded by Osaka fibrils replicated the Osaka fibril structure in WT Aβ_1-40_ monomer. Specifically, when Osaka fibril seeds were incubated with WT Aβ_1-40_ monomer, the two main morphologies that were produced were a 50 nm crossover distance (40 ± 26%) and no crossovers (23 ± 10%). Additional morphologies were observed when Osaka fibril seeds were incubated with WT Aβ_1-40_ monomer including 40 nm (16 ± 15%), 60 nm (8 ± 1%), 120 nm (5 ± 5%), and 200 nm (2 ± 2%).

To evaluate if this seeding effect is reciprocal between mutants and WT Aβ_1-40_, we also examined the effect that WT Aβ_1-40_ fibril seeds have on monomeric Aβ mutants. As seen in Figure 2c and 2d, when Arctic and Osaka monomers were seeded by WT Aβ_1-40_ fibrils, their fibril population distribution remained similar to that when aggregated in the bulk solution. In the case of WT Aβ_1-40_ fibrils seeding Arctic mutants, the resulting fibril morphologies largely consisted of crossover distances of 40 nm (58 ± 30%) and 30 nm (34 ± 29%). Similarly, WT Aβ_1-40_ fibril seeds had little effect on the formation of Osaka fibrils, as the primary fibril morphology produced had a crossover distance of 50 nm (66 ± 5%).

In contrast to the results observed when cross-seeding WT Aβ_1-40_ with mutants, cross-seeding WT Aβ_1-40_ with its isoform WT Aβ_1-42_ yielded a bimodal distribution of crossover distances for one combination and complete heterogeneity for the other. When WT Aβ_1-42_ was incubated alone, it primarily formed fibrils with an 18 nm crossover distance (64 ± 24%), a structure that was not seen in WT Aβ_1-40_ (Figure 2e). However, when WT Aβ_1-42_ monomer was incubated with WT Aβ_1-40_ fibrils, a bimodal distribution appeared; the proportion of fibrils with a crossover distance of 18 nm was approximately half (38 ± 3%) of what was observed in the WT Aβ_1-42_ only condition, and the rest of the fibril structural distribution more closely resembled the distribution observed in the WT Aβ_1-40_ only condition. When comparing the structural distributions of WT Aβ_1-40_ or WT Aβ_1-42_ seeded by the Osaka fibrils, we saw that while WT Aβ_1-40_ took on a similar structural distribution to that of the Osaka, the WT Aβ_1-42_ developed a similar bimodal distribution to that observed when WT Aβ_1-42_ monomer was seeded with WT Aβ_1-40_ fibril, shown in Figure S1. The two dominant structures were either the 18 nm crossover distance fibrils seen from WT Aβ_1-42_ fibrils alone (38 ± 12%) and the 50 nm crossover distance fibrils (33 ± 9%) seen from Osaka mutant fibrils alone (Figure 2e). However, when WT Aβ_1-40_ monomer was incubated with WT Aβ_1-42_ fibrils, the resulting WT Aβ_1-40_ fibrils had a wide distribution of fibril structures, ranging in crossover distance from 50 nm to 250 nm, as well as a large population of fibrils with no crossovers.

### Kinetic Analysis of Mutant Fibril Formation

To further probe the differences between Arctic, Osaka, and WT Aβ_1-40,_ we also measured the rate of fibril formation by monitoring Thioflavin T (ThT) fluorescence of 30 μM Aβ_1-40_ incubated at 37 °C in a 10 mM sodium phosphate (NaPO_4_) buffer. In this assay, fluorescence intensity correlates with fibril formation as ThT intercalates in the aggregate’s increasing β-sheet content. As seen in Figure 3, similar to the structural seeding experiments, 5 μM of pre-formed Arctic (Figure 3a) and Osaka (Figure 3b) fibrils were able to seed the fibrilization of 30 μM monomeric WT Aβ_1-40_, shown by the increase in fluorescence, in the case of the Arctic mutation, and the decreased lag phase, in the case of the Osaka mutation. This kinetic result is consistent with our structural analysis in which pre-formed Arctic and Osaka fibrils pass on their structure into WT Aβ_1-40_. With each subsequent generation of WT Aβ_1-40_ monomer introduced to the previous generation’s fibrils, the aggregation kinetics further accelerated as indicated by the subsequent decrease in lag phase (Figure 3e). Similar to previous literature reports,^46^ monomeric WT Aβ_1-40_ added to the Arctic mutation formed fibrils almost instantaneously (Figure 3c). Additionally, Aβ_1-40_ with the Osaka mutation did not interact with ThT until later times (Figure 3d), consistent with previous reports.^55^

**Figure 3.**
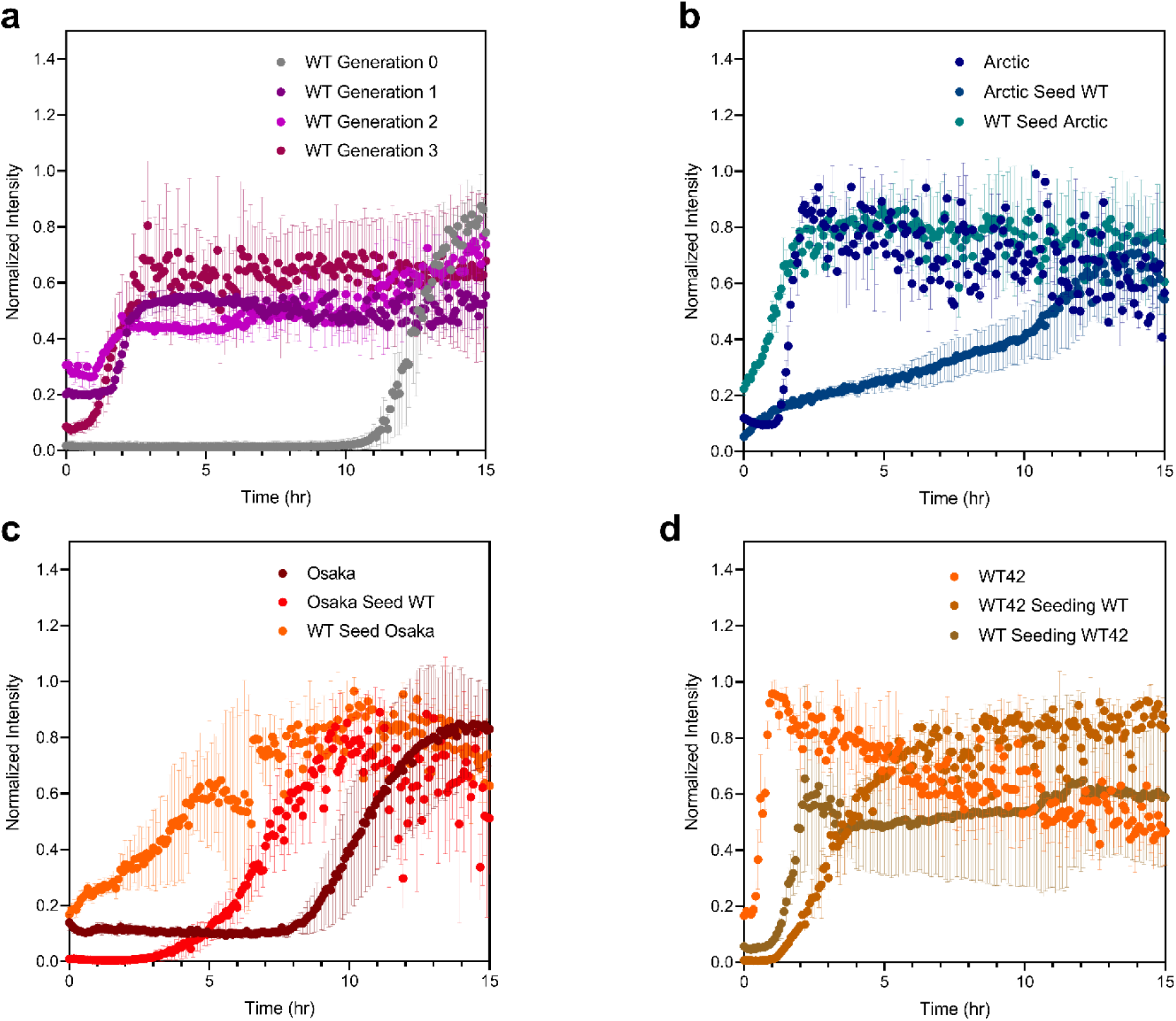
Fluorescence curves of the kinetics of aggregation for (a) 30 μM WT (grey) and 5 μM Arctic seeding 30 μM WT (blue), (b) 30 μM WT (grey) and 5 μM Osaka seeding 30 μM WT (red), (c) 30 μM Arctic (dark blue) and 5 μM WT seeding 30 μM Arctic (light blue), (d) 30 μM Osaka (dark red) and 5 μM WT seeding 30 μM Osaka (orange), and (e) Generations 0 to 3 of 5 μM WT seeding 30 μM WT. All samples were run at 37°C with 100 μM thioflavin T. Error bars represent one standard deviation from three separate experiments.

Interestingly, we observed similar accelerated kinetics when monomeric Aβ_1-40_ with the Arctic and Osaka mutation was seeded by pre-formed WT Aβ_1-40_ fibrils (Figures 3c and 3d). In this case, ThT fluorescence was immediately detected, indicating that WT fibrils rapidly seeded the formation of fibrils for the mutant Aβ_1-40_. However, we did not observe the same structural effect when analyzing the crossover distances of Arctic and Osaka fibrils seeded by WT. As seen in Figure 2c and 2d, the distribution resembled that of the mutants formed in the absence of seeds. Therefore, it is possible that the WT seeds are providing a nucleating surface for the Arctic and Osaka fibrils to aggregate more quickly, but the WT seeds are not replicating their structure into fibrils with the Arctic and Osaka mutation.

For WT Aβ_1-42_ and cross-seeding both ways with WT Aβ_1-40_, we observed a very small difference in aggregation kinetics. Pure WT Aβ_1-42_ had a lag time that was less than one hour, whereas cross-seeding both directions with WT Aβ_1-40_ led to lag times of roughly one hour. With the lag times so similarly quick across all three conditions, it is clear cross-seeding WT Aβ_1-40_ monomer with WT Aβ_1-42_ fibril seeds accelerate kinetics. However, cross-seeding WT Aβ_1-42_ monomer with WT Aβ_1-40_ fibril seeds has little effect on aggregation kinetics.

### Analysis of Cell Viability Phenotypes from Mutant Fibrils

To further probe the phenotypes of the various structures observed in WT and mutant fibrils, we measured their effect on cellular health. We added 30 μM of the Aβ fibrils to cultures of SH-SY5Y human neuroblastoma cells and measured cell viability using a assay based on the absorbance of (3-(4,5-dimethylthiazol-2-yl)-2,5-diphenyltetrazolium bromide (MTT) converted to the reduced formazan product in the presence of viable cells. This assay provides a quantitative assessment of cellular activity as a proxy for *in vivo* toxicity of amyloid fibrils. We first measured the viability of SH-SY5Y cells after a 24 hr incubation with Generation 0, 1, 2, and 3 WT Aβ_1-40_ fibrils (Figure 4a) to determine the effect of fibril crossover distance on cell health. Generations 0 and 1 had a cell viability of 78 ± 6% and 97 ± 20% compared to untreated cells, respectively, which then dropped in Generations 2 and 3 to 75 ± 6% and 54 ± 6%, respectively. As crossover distance decreased across generations, fibrils showed more toxicity to these neuroblastoma cells (Figure 4a).

**Figure 4.**
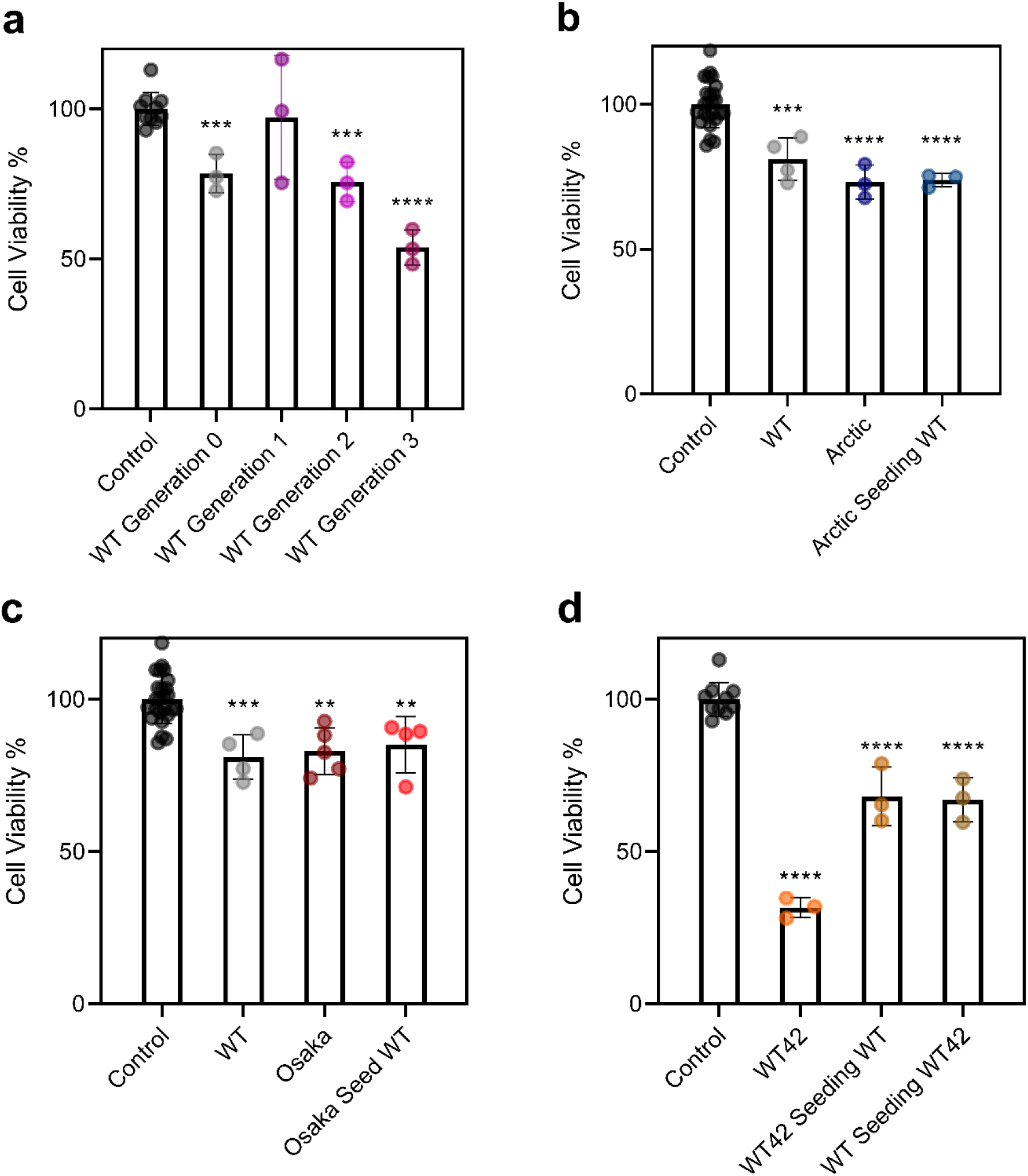
The cytotoxic effects of (a) repeated seeds of Aβ_1-40_; (b) mutant Arctic fibrils and their WT cross-seeds repeated seeds of Aβ_1-40_; (c) mutant Osaka fibrils and their WT cross-seeds; (d) and WT Aβ_1-42_ fibrils and their WT Aβ_1-40_ cross-seeds on SH-SY5Y neuroblastoma cells. Cytotoxicity was measured using an MTT assay after adding 30 μM fibrils to cells for 24 hr. Error bars represent one standard deviation from three independent measurements. *,**,***, and **** represent significant differences as measured by a one-way ANOVA (* = p < .05), (** = p < .01), (*** = p < .001), and (**** = p < .0001).

Next, we analyzed the effects of E22 mutant fibrils and their respective seeds on cell health. While all the fibrils displayed a statistically significant decrease in cell viability (Figure 4b and 4c), the results show that Arctic fibrils were slightly more toxic (73 ± 6%) than Osaka (82 ± 7%) and WT (81 ± 7%) fibrils. Additionally, the progeny WT fibrils formed when seeded by Arctic and Osaka fibrils displayed a similar toxicity profile to their parent seeds. Specifically, Arctic seeding WT fibrils had a cell viability of 73 ± 2%, while Osaka seeding WT fibrils had a cell viability of 85 ± 9%.

As an additional comparison, we also examined the cytotoxicity of Aβ_1-42_ alone and cross-seeded with WT Aβ_1-40_ The WT Aβ1−42 isoform, which is known to be more toxic than WT Aβ_1-40_, produced a unique fibril structural distribution (Figure 2e) with a crossover distance primarily of 18 nm (64 ± 24 %). WT Aβ_1-42_ displayed the lowest viability (32 ± 3%) out of all the fibrils. When WT Aβ_1-42_ was cross-seeded by WT Aβ_1-40_ fibrils, the viability was between the observed results of both WT Aβ_1-40_ alone and WT Aβ_1-42_ alone (67 ± 7%) (Figure 4d). A similar cellular viability was also observed when WT Aβ_1-40_ was cross-seeded by WT Aβ_1-42_ fibrils (68 ± 10%).

Overall, we observed that regardless of monomer sequence, cellular viability closely correlated with crossover distance. Short fibril crossover distances like those observed in the WT Aβ_1-40_ generation 3, Arctic, and WT Aβ_1-42_ lead to poorer cellular health outcomes compared to the other fibril distributions with longer crossover distances. These results also indicate that when mutant fibrils, such as Arctic, replicate their helical crossover into WT fibrils, they confer similar cell toxicity phenotypes into WT fibrils, implying that this phenotype *in vivo* may depend on structure and not sequence. Indeed a similar effect has been observed in transgenic mice expressing WT Aβ_1-40_ that were injected with Arctic fibrils, after which the mice began showing hallmarks of familial Alzheimer’s disease (FAD).^56^

In addition to measuring cell viability as a metric for cell health through the MTT assay, we also evaluated anilinonapthalene-8-sulfonate (ANS) binding for differences in fibril surface hydrophobicity, as well as the effect of the mutant fibrils and their respective seeds on the generation of reactive oxygen species (ROS) in SH-SY5Y cells. For ANS binding, we found that WT fibrils maintained similar hydrophobicity regardless of seeding conditions, whereas both mutant fibrils had increased surface hydrophobicity, as seen in Figure S2. Oxidative stress from ROS generated both intra- and extracellularly is a hallmark characteristic of AD; measuring ROS generation may therefore provide insight into the Aβ variants’ role in disease pathology.^57^ To measure the generation of ROS, we used 2’7’-dichlorofluorescein diacetate (DCFH-DA), which fluoresces strongly when excited in the presence of ROS. Compared to untreated SH-SY5Y cells, both Arctic and Osaka fibrils displayed a statistically significant increase in ROS production (Figure S3). Conversely, WT Aβ fibrils and WT fibrils seeded by the mutants demonstrated only a small increase in ROS production when compared to the untreated cells. This suggests that similar to the ANS binding assays, this phenotype is potentially controlled by sequence instead of structure, as opposed to fibril cellular toxicity when measured with an MTT assay. This may also suggest that ROS is not what is killing these cells. The polymorphs from Osaka and Arctic increased ROS and to understand better this observation, we will need higher resolutions structures. This is a topic of ongoing work, in our laboratory and others.

## Discussion

The objective of this work was to investigate how the cross-seeding of different Aβ sequences influences the distribution of amyloid fibril structures and their resulting phenotype. Previous studies of fibril cross-seeding have recognized that monomers or aggregates of different sequences can interact and reciprocally induce fibril formation during the progression of amyloid-associated diseases.^40^ These studies primarily investigated changes in aggregation kinetics using ThT to determine if different protein sequences could interact during fibrillization.^39^ In contrast, we examined cross-seeding induced structural and phenotypic changes to the entire fibril population through a combination of EM analysis, kinetics, and cellular assays. Specifically, we monitored changes in the entire population distribution of fibril polymorphs from cross-seeding to better understand the complexity of samples generated *in vivo*, and to determine whether the resulting functions of these fibril structures were changing in ways that could be attributed to one specific distribution of polymorphs, or even one dominant member of the population.

We initially investigated the distribution of fibril polymorphs, which were characterized by different crossover distances, formed through repeated seeding of WT Aβ_1-40_. Initial fibrilization (Generation 0) of this sequence formed a highly heterogenous population of fibrils with a variety of crossover distances, with particularly high concentrations found at 60 nm, 120 nm, or no measurable crossover. Upon repeated seeding with fresh Aβ_1-40_ monomer, the fibril population became more homogeneous and was dominated by a population of approximately 30 nm crossover distance fibrils. Notably, our results are consistent with previous studies where repeated seeding was used to prepare fibril populations suitable for solid-state NMR (ssNMR) spectroscopy.^22,41^ Our results also demonstrate that fibril structures that were almost entirely absent in the initial aggregation event can nevertheless dominate the population under the right conditions. For example, it was recently found that for WT Aβ_1-42_, small fibril populations could dominate the structural seeding through secondary nucleation of additional monomer under some conditions.^58^ Ultimately, repeated seeding of the same amyloid-forming monomer may be an important tool in forming homogeneous fibril populations or replicating *ex vivo* structures under *in vitro* conditions.

Aβ mutants and isoforms associated with increased disease pathologies are known to exhibit unique fibril structure characteristics.^36^ However, the simultaneous presence of multiple isoforms (Aβ_1-40_ and Aβ_1-42_) or sequences (WT, Arctic, and Osaka) in individual patients complicates the fibrillization landscape and makes understanding cross-seeding relationships highly physiologically relevant. Thus, we investigated the effects of cross-seeding on fibril structural populations for FAD mutants (Arctic and Osaka), Aβ_1-40_, and Aβ_1-42_. The first generation of fibrils of each sequence investigated, WT Aβ_1-40_, WT Aβ_1-42_, Arctic, and Osaka, resulted in a heterogeneous population of fibril structures that had crossover distances ranging from 30 nm to 200 nm, as well as fibrils with no observed crossovers. Importantly, the pattern of dominate observed crossover distances was distinct for each sequence, and indeed could even be used as a fingerprint of the specific mutation. Next, we used these structures to seed one generation of WT Aβ_1-40_ fibrils; these structures, as seen in Figure 2c and 2d, can be compared directly to Generation 1 of WT Aβ_1-40_ self-seeding (Figure 2b). WT Aβ_1-40_ fibrils seeded with either Arctic or Osaka fibrils had homogeneous crossover polymorph distributions that were highly similar to the Generation 0 Arctic and Osaka fibrils. In other words, the Arctic and Osaka sequences imprinted their aggregated fibril structure onto the WT Aβ_1-40_ sequence, resulting in dramatically different fibril structures. However, the opposite was not true; Arctic and Osaka Aβ monomers produced the same fibril structures regardless of whether seeds of WT Aβ_1-40_ fibrils were present. Our results suggest that fibrils with shorter crossover distances may be more effective at structurally templating new fibrils, regardless of sequence similarity, apart from the dominant WT Aβ_1-42_ fibril with an 18 nm crossover distance. Although the exact mechanism is unclear, this possibility is also supported by repeated isolation and structural characterization of shorter crossover distance fibrils from disease patients.^32,33,59–61^ The ability of mutants/isoforms to bias the structural population of Aβ_1-40_ fibrils into potentially more toxic conformations may also have important disease implications.^59^

Our cross-seeding results with WT Aβ_1-40_ and WT Aβ_1-42_ were consistent with previous studies, where seeding WT Aβ_1-40_ monomer with WT Aβ_1-42_ fibrils was inefficient at producing the same fibril structure of WT Aβ_1-42_.^33^ There was some success, however, when we cross-seeded WT Aβ_1-42_ monomer with WT Aβ_1-40_ fibrils. We attribute this to the asymmetric surface that some polymorphs of WT Aβ_1-42_ fibrils display compared to the symmetric surface WT Aβ_1-40_ and its mutants display, as shown in ssNMR studies.^36^ Differences in cross-seeding efficiency were also observed in previous repeated seeding work using type II diabetes patient-derived human islet amyloid polypeptide (hIAPP) fibrils. In one study, roughly 10% of the resulting seeded hIAPP fibrils were nearly identical to the pathogenic seeds, whereas most of the seeded polymorphs differed from the unseeded controls.^59^ A notable finding in our experiments was that in both cases where WT Aβ_1-42_ monomer was seeded by a fibril that was either WT or Osaka Aβ_1-40_, the resulting distribution was bimodal, where half the fibrils were similar to pure WT Aβ_1-42_ fibrils and half of the fibrils were similar to whichever parent fibril was used to seed the monomer.

Next, we measured the effect of successive generations of seeding and cross-seeding on the kinetics of fibril formation. As expected, the presence of seeds from any sequence accelerated fibrillization kinetics, except for WT Aβ_1-40_ fibrils seeding WT Aβ_1-42_ monomer, which had a very minimal effect. However, faster aggregation kinetics did not necessarily lead to shorter crossover distances. For example, WT Aβ_1-40_ fibril seeds accelerated the aggregation of fresh Aβ_1-40_ monomer but did not appreciably affect the distribution of different fibril structures until seeding generation 2. The independence of structure and aggregation kinetics was especially noticeable in fibrils formed from Arctic and Osaka monomers, which retained their structure in the presence of WT Aβ_1-40_ seeds, despite aggregating faster than the mutant Aβ_1-40_ monomer alone. The complex set of structural and kinetic interactions between the four Aβ sequences investigated here are summarized schematically in Figure 5. The concentration or number of seeds can significantly influence the predominance of primary or secondary nucleation during fibril formation.^58^ While we only examined a single seed concentration, an exciting complement to previous kinetic studies may be examining how fibril structural populations change as a function of seed concentration. Collectively, our data highlight the complex interplay between aggregation kinetics and resulting fibril structure while also indicating that faster kinetics may not necessarily indicate the formation of a specific fibril structure.

**Figure 5.**
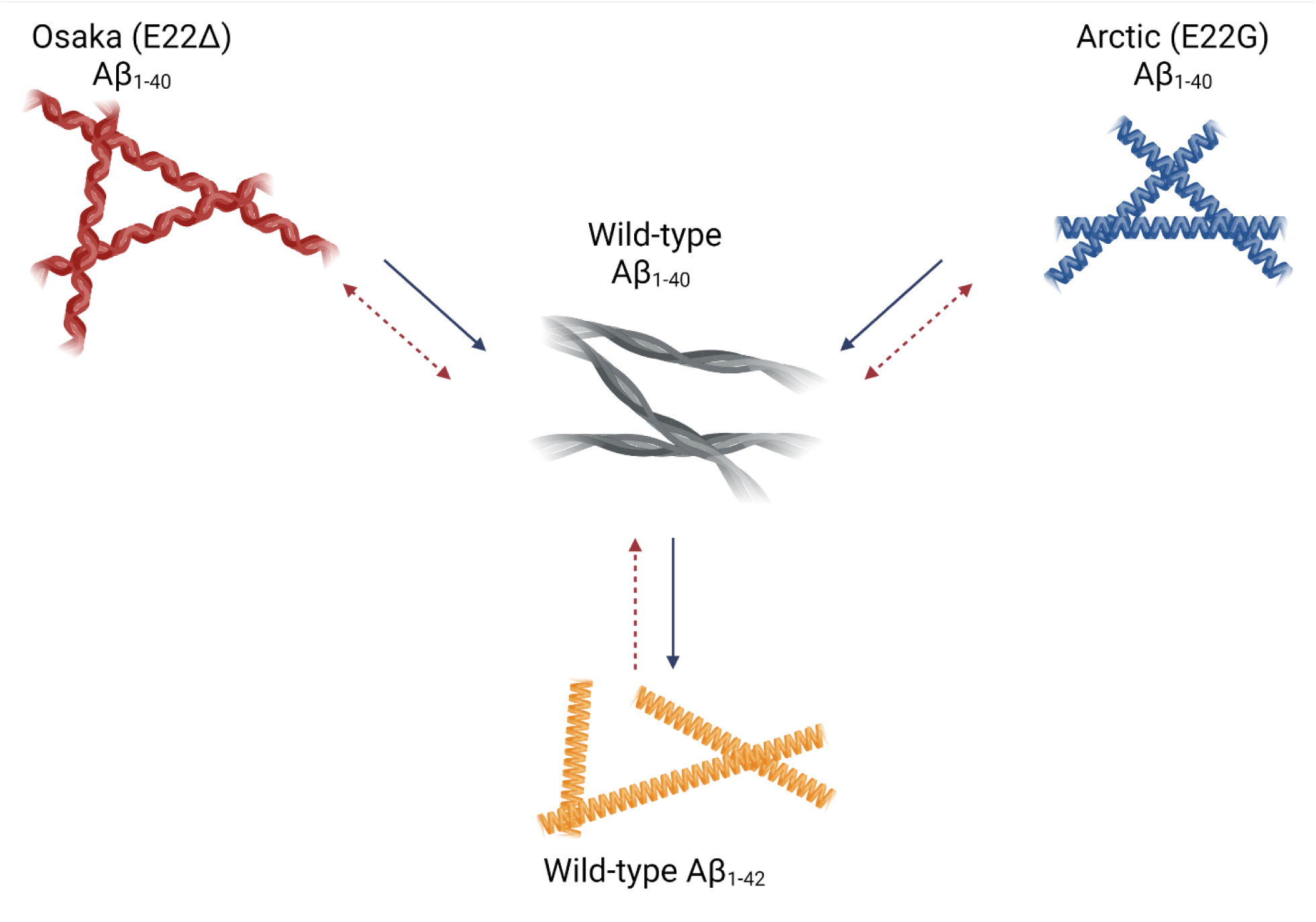
Summary schematic of cross-seeding of Aβ mutants and isoforms. Solid blue arrows represent the ability to seed fibril structure of the parent sequence. Dashed red arrows represent the ability to affect the kinetics of fibril formation.

Finally, we investigated the effect of different fibril populations on cell health. Even with relatively coarse cytotoxicity assays, we measured significant differences in cytotoxicity between Aβ_1-40_ seeded at different generations and with the Aβ_1-40_ mutants. Generally, shorter crossover distances fibrils were more cytotoxic. For example, repeated seeding of WT Aβ_1-40_ generated higher populations of short crossover fibrils, which showed higher toxicity. Consistent with literature reports, WT Aβ_1-42_ fibrils were comprised of mainly shorter crossover distance structures and showed higher cytotoxicity than Aβ_1-40_. However, WT Aβ_1-42_ toxicity was attenuated when it was seeded using WT Aβ_1-40_ fibrils, possibly due to a higher percentage of longer crossover fibrils. WT Aβ_1-40_ toxicity slightly increased when it was seeded using WT Aβ_1-42_ fibrils despite retaining similar heterogeneity to WT Aβ_1-40_ fibrils incubated alone, warranting further investigation on the impact WT Aβ_1-42_ fibrils have when seeding WT Aβ_1-40_ monomer. Crossover distance was not the sole determinant of cytotoxicity as both Arctic and Osaka monomers formed high populations of fibrils with 50 nm or shorter crossover distances but had relatively minimal effects on viability. We note our results are broadly consistent with recent high resolution cryo-EM structures of fibrils isolated from disease patients, where significant proportions of presumably toxic fibril polymorphs with shorter crossover distances were identified. For Aβ fibrils from Alzheimer’s patients, 28% of the fibrils had a crossover distance of roughly 50 nm.^32^ In multiple system atrophy patients, all of the polymorphs observed had crossover distances of roughly 60 nm.^60^ For type II diabetes, observed polymorphs of hIAPP fibrils seeded from patient-derived fibrils successfully demonstrated toxicity had crossover distances of roughly 26 or 60 nm.^43^ The exact mechanism for how fibril sequence, structure, and polymorph distribution combine to influence cellular health remains unknown. One possibility of shorter crossover distance leading to higher toxicity could be due the more frequent number of changes to the fibril’s chemical profile across a fixed distance of the cell membrane. Because Aβ in its monomeric and oligomeric forms is observed to have a detergent-like quality that disrupts membrane lipids due its amphiphilic nature,^62^ shorter crossover distances may enable more frequent changes of the fibril’s polar and non-polar amino acid interactions with the membrane along the fibril axis, which could further accentuate this detergent-like quality.^61^ Ultimately, while *in vitro* fibril preparations can successfully recapitulate more complex neurodegenerative disease phenotypes,^59^ we urge caution in interpreting these results since our data indicate that fibril structural populations and biochemical properties can change dramatically.

In conclusion we measured the population distribution and biochemical properties of Aβ fibrils formed through repeated self- and cross-seeding. Our work provides critical information on the complex relationship between fibril structure(s) and function, which could be leveraged toward a better understanding of neurodegenerative disease pathology or the preparation of *in vitro* fibrils that are more structurally representative of disease isolates. The ability of Aβ_1-40_, and presumably other amyloid forming proteins to rapidly adopt new structures after repeated seeding may also explain patient-to-patient variation, amyloid drug resistance,^63^ or selection of beneficial amyloid phenotypes.^64,65^ Finally, our work provides insight into observations of how mixed genetic variants of WT Aβ_1-40_ sequences can lead to unique differences in disease pathology. For example, mice that are heterozygous carriers of the Osaka mutation do not display signs of dementia, while those while those with the Arctic mutation do.^47,48^ Future work should focus on the effects of other common Aβ isoforms and post-translational modifications on the properties and phenotypes of the resulting fibril populations. Overall, supplementing high resolution fibril structures with snapshots of heterogeneous fibril populations is a significant step towards understanding amyloid biophysics and neurodegenerative disease pathology.

## Materials and Methods

### Preparation of Aβ_1-40_

5 mg of Aβ_1-40_ (BACHEM, H-1994), 1 mg of Arctic Aβ_1-40_ (BACHEM, 4035372), and 1 mg of Osaka Aβ_1-40_ (BACHEM, 4091431) were dissolved in 1mL, 500 μL, and 500 μL of hexafluoro-2-propanol (HFIP) and shaken at 300 RPM for 30 min. 40 μL aliquots were then added into microcentrifuge tubes, and the HFIP was allowed to evaporate overnight. The dried peptide samples were then further dried through vacuum centrifugation for 30 min. The dried Aβ samples were then stored at −20 °C.

### Fibrilization of Aβ_1-40_ and Mutants for Structural Analysis

An aliquot of monomer was thawed and then dissolved in 1% dimethyl sulfoxide (DMSO) and 10 mM sodium phosphate (NaPi) buffer to produce a 60 μM solution of Aβ_1-40_. Aliquots (100 μL) of dissolved monomer were then pipetted into a non-binding, coated 96-well plate (Corning 3991). Each well was then diluted with an additional 100 μL of NaPi to produce a final volume of 200 μL and a Aβ_1-40_ concentration of 30 μM. The plate was sealed and incubated at 37 °C, shaking at 300 RPM. Each sample was run in triplicate for structural analysis. Fibril “seeds” for different peptide monomers that were used for additional seeding experiments were prepared using the same workflow.

### Transmission Electron Microscopy

Transmission electron microscopy was used to analyze the morphology of the resulting Aβ fibrils. Samples were prepared on carbon coated grids (Electron Microscopy Sciences, CF400) that were glow discharged with an EmiTech K100x Coater. After the amyloid samples were allowed to aggregate for 24 hr, 7 μL of the amyloid suspension was drop-cast on the charged carbon side of the grid. After incubating for 1 min, the droplets were washed two times each with in 50 μL of 0.1 and 0.01 M ammonium acetate. The droplet was then washed with 50 μL of 2% uranyl acetate. Following the washes, excess liquid was wicked away using filter paper. Negative-stain images were acquired on a JEOL 2010F TEM operated at 200 kV at a nominal magnification of 60000× and a JEOL NeoARM operated at 200 kV, both settings of which resulted in a pixel size of 3.6 Å/pixel. The exposure for each image was 2 s, resulting in a total electron dose of 60–70 e– Å–2. Data was collected on Gatan OneView cameras with a defocus ranging from ™1.0 to ™3.5 μm.

### Image Processing

Fibril segments were picked using eman2helixboxer from the EMAN2^66^ image processing suite, with a box size of 556 pixels (200 nm), no overlap, and one side of the box always selected at the start of a fibril crossover. This box size was selected based on the measurement of the longest helical crossover distance observed. Fibril particles were manually picked using EMAN2 from 3 biological replicates that each contained 50 micrographs, resulting in a total of 150 images per condition. These were then extracted and segmented into particles with a box size of 1238.4 Å using RELION. Approximately 1,950 particles were used to measure the distributions of fibrils with different helical crossover distances across conditions. The particles were further refined by 2D classification into three classes using RELION with helical symmetry optimization enabled.

### Kinetic Assays of Aβ_1-40_ and Mutant Aggregation

For both pure WT and mutant Aβ_1-40_, samples were prepared in a similar fashion as the preparation for structural analysis. Replicates were prepared in a non-binding 96-well plate at a final volume of 200 μL and a concentration of 30 μM Aβ_1-40_. For seeding experiments, wells were prepared with a final concentration of 5 μM fibril seeds and 30 μM monomer. Additionally, 2 μL of 10 mM thioflavin T (ThT) was added to each well for a final concentration of 100 μM ThT. The plate was sealed with spectroscopy grade tape (Thermo Scientific 235307), and fluorescent measurements were taken using a fluorescent plate reader (Clariostar, BMG Labtech) at 37 °C. ThT fluorescence was measured with an excitation wavelength of 440 nm and an emission wavelength of 480 nm through the top of the plate. Measurements were taken every 300 sec for 48 hr. Prior to each measurement, the plate was shaken at 300 RPM. Each sample was run in triplicate with blanks (without Aβ_1-40_) to account for background ThT fluorescence Blanks with seeds were also prepared to account for ThT interaction with the 5 μM fibril seeds.

### Preparation of Peptide for Cellular Assays

A 100 μL aliquot of each sample was centrifuged at 21,000 RCF for 10 min, the supernatant was removed, and the pellet was suspended in 50 μL HFIP. The HFIP was allowed to evaporate overnight and then further dried by vacuum centrifugation. The dried peptide was resuspended in 100 μL 10 mM NaPi and the concentration was estimated using a Bradford assay, in which 200 μL of Coomassie Plus (Thermo Scientific 23236) was mixed with 20 μL of sample. Bovine serum albumin (BSA) was used as a standard for a calibration curve. Once concentration of fibril was determined, the original sample was centrifuged at 21,000 RCF for 10 min, the supernatant was removed, and the pellet was resuspended in DMEM without phenol red, glucose, or glutamine to bring the final concentration to 30 μM.

### Cytotoxicity Assays

SH-SY5Y human neuroblastoma cells (ATCC CRL-2266) were cultured in a 1:1 media mixture of EMEM:F12 with 10% FBS and 100 U mL^-1^ penicillin-streptomycin at 37 °C in a 5% CO_2_ environment. Cells were used for assays after the 6^th^ passage and before the 10^th^ passage.

The cells were plated at 4 × 10^4^ cells well^-1^ in a Nunclon Delta-Treated 96-well plate (Thermo Scientific 167008). After allowing an overnight growth for cells to adhere, the plates were centrifuges at 500 RCF for 5 min, aspirated, and washed with phosphate buffered saline (PBS). Then, 200 μL of 30 μM Aβ_1-40_ fibrils suspended in Dulbecco’s modified eagle’s medium (DMEM) without phenol red, glucose, and glutamine, was added to each well. All samples were run in at least triplicate. Control wells contained only 200 μL of DMEM without any fibrils. After a 24 hr incubation, the plates were centrifuged and aspirated. 100 μL of DMEM and 10 μL of CyQUANT MTT Cell Viability Assay in PBS (Thermo Scientific V13154) was then added. After a 4 hr incubation with the MTT, cells were centrifuged, aspirated, and resuspended in 100 μL of dimethyl sulfoxide (DMSO). Absorption measurements were collected at 490 nm using a plate reader (Clariostar, BMG Labtech).

### Reactive Oxygen Species Assays

SH-SY5Y neuroblastoma cells were cultured as described above. Cells were plated at 4 × 10^4^ cells well^-1^ in a Nunc Microwell, Optical Polymer Base 96-well plate (Thermo Scientific 165305) and allowed to grow overnight. Cells were then washed with 100 μL of PBS and incubated with 100 μL of 20 μM 2’,7’-dichlorofluorescin diacetate (DCFH-DA, Sigma D6883) in DMEM in the dark for 40 min at 37 °C. Cells were then washed with 100 μL of PBS and treated with 100 μL of 30 μM of Aβ_1-40_ in DMEM without phenol red, glucose, or glutamine. Control wells were prepared without and DCFH-DA and without peptide. Additionally, cells were treated with 5 mM *t*-butyl hydroperoxide (*t*-BHP) as a positive control, representing 100% ROS production. After a 6 hr incubation, fluorescence intensity was collected using a plate reader (Clariostar, BMG Labtech), with an excitation wavelength of 480 nm and an emission wavelength of 535 nm.

### ANS Binding Assay

Using a Greiner 96-well plate, 200 μL samples were prepared in triplicate with 7.5 μM of WT or mutant Aβ_1-40_ in 10 mM NaPi. Aliquots of 3 μL of 5 mM 8-anilino-1-naphthalene sulfonic acid (ANS) were added to each well. The fibril samples were incubated with ANS in the dark at 37 °C for 10 min, and then analyzed for fluorescence in a plate reader (Clariostar, BMG Labtech). Emission spectra from 400-600 nm were collected using an excitation wavelength of 370 nm.

### Statistical Analysis

Unless otherwise noted, data are reported as a mean ± S.D. of *n* = 3 biological replicates. Significance for cellular assays was calculated in GraphPad Prism 8.0 using a one-way ANOVA (α = 0.05).

## Acknowledgements

This work was supported in part by Welch Foundation Research Grants F-1722 (to L.J.W.) and F-1929 (to B.K.K.); National Science Foundation CHE-1807215 (to L.J.W.); and the College of Natural Sciences at The University of Texas at Austin (Catalyst Grant to L.J.W., B.K.K., and D.W.T.). Partial support was provided by the National Science Foundation through the Center for Dynamics and Control of Materials: an NSF Materials Research Science and Engineering Center under DMR-1720595. D.W.T. is a CPRIT Scholar supported by the Cancer Prevention and Research Institute of Texas (RR160088) and an Army Young Investigator supported by the Army Research Office (W911NF-19-1-0021). Additionally, we are grateful for the use of the facilities within the Texas Materials Institute and the technical expertise provided by Dr. Karalee Jarvis.

