## Supplemental Figures for "Cross-seeding Controls Aβ Fibril Populations and Resulting Function"

### Supporting Information

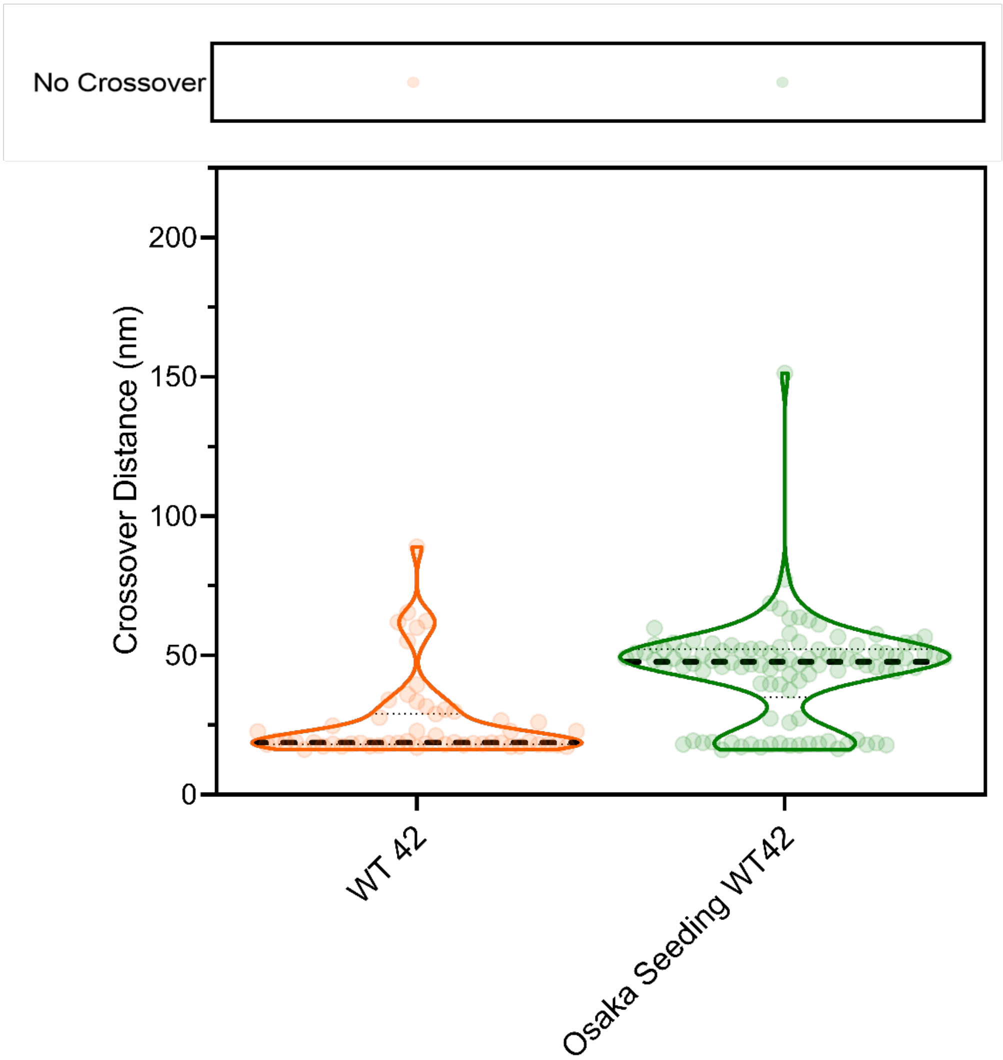


Figure S1. Distribution of fibril crossover distances for A$\beta_{1\text{-}42}$ fibrils and 5 μM Osaka seeding A$\beta_{1\text{-}42}$ fibrils at 37°C. 50 micrographs were analyzed for each of three biological replicates on negative-stain EM. Dashed line represents the median.

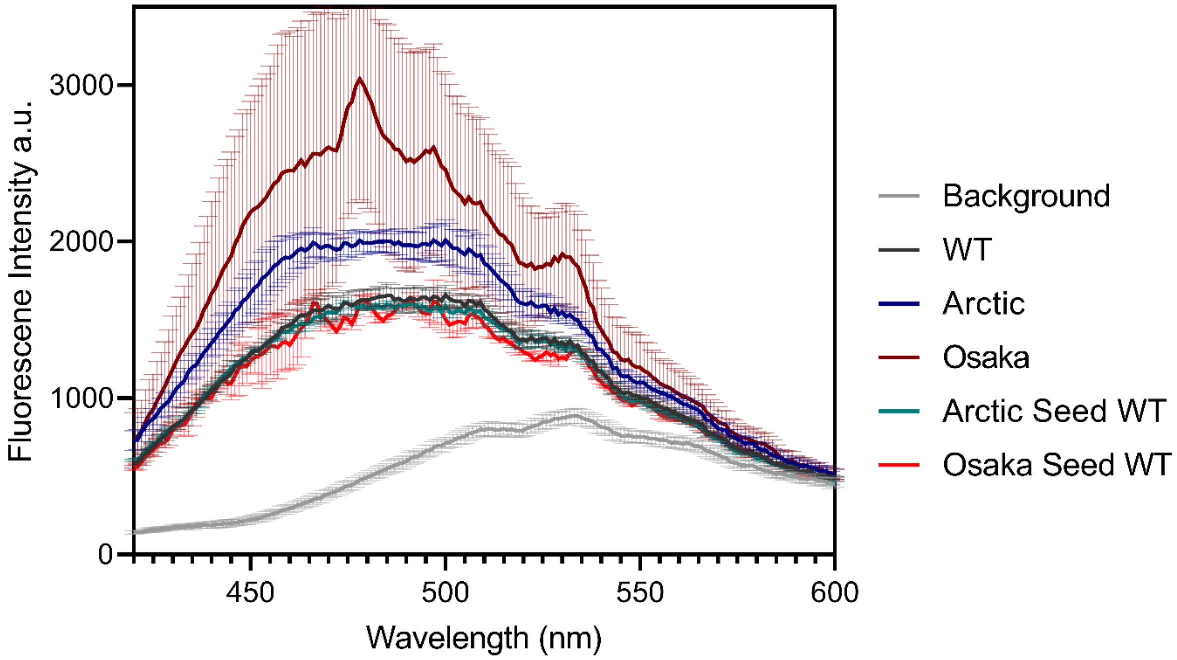


Figure S2. ANS Binding fluorescence spectra of 7.5 µM WT A$\beta_{1\text{-}40}$ fibrils, mutant A$\beta$ fibrils, and fibrils of A$\beta_{1\text{-}40}$ monomer cross-seeded with mutant A$\beta$ fibril seeds in 10 mM NaPi, incubated with ANS in the dark at 37 °C for 10 min in triplicate.

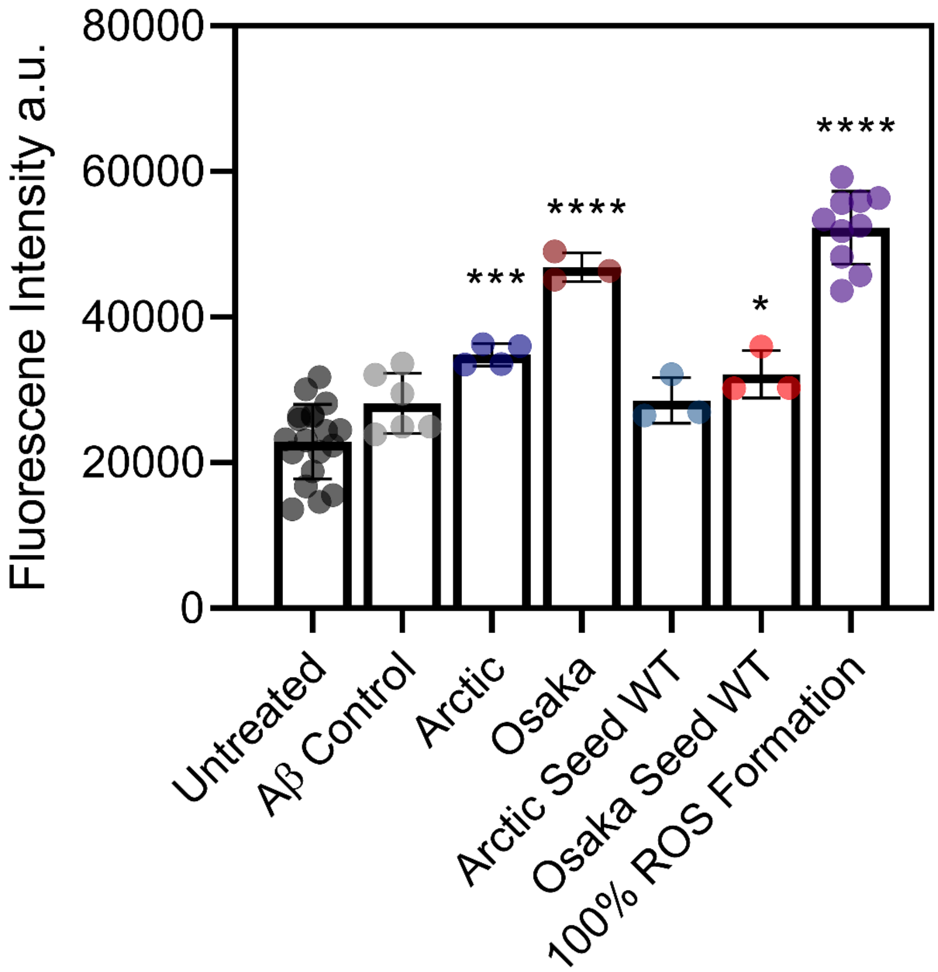


Figure S3. The production of reactive oxygen species (ROS) induced by 30 µM A$\beta_{1\text{-}40}$ mutants and WT seeds in SH-SY5Y neuroblastoma cells after 6 hr. ROS production was measured using a DCFH-DA assay. For the 100% ROS control, cells were treated with 5 mM *t*-BHP. Error bars represent 1 SD. *,**,***, and **** represent significant differences as measured by a one-way ANOVA (* = $p < .05$), (** = $p < .01$), (*** = $p < .001$), and (**** = $p < .0001$)
